# *Cladosporium herbarum* peptidogalactomannan triggers significant defense responses in whole tobacco plants

**DOI:** 10.1101/2020.11.12.379552

**Authors:** Caroline de B. Montebianco, Bianca B. Mattos, Tatiane da F Silva, Eliana Barreto-Bergter, Maite F S Vaslin

## Abstract

*Cladosporium herbarum* is one of the most frequently occurring fungal species, with a worldwide distribution, and is found in almost all man-occupied niches in organic and inorganic matter and as a phytopathogen on certain agricultural crops. The structure of the most abundant glycoprotein from the *C. herbarum* cell wall, peptidogalactomanann or pGM, was previously elucidated and includes carbohydrates (76%), with mannose, galactose and glucose as its main monosaccharides (52:36:12 molar ratio). pGM was able to strongly induce the expression of defense-related genes and ROS accumulation when in contact with BY2 tobacco cells. Here, using two distinct *Nicotiana tabacum* cultivars, Xanthi and SR1, we evaluated the ability of *C. herbarum* pGM to induce SAR-like defense by studying its antiviral activity against *Tobacco mosaic virus* (TMV) and the induction of SAR markers including *PR* genes and ROS accumulation. Our results show that pGM induced a strong activation of defense responses in treated plants from both tobacco cultivars, contributing to the impairment of viral infection. Expression levels of the pathogenesis-related genes *PR-1a* (unknown function), *PR-2* (□-1-3 *endoglucanase*) *PR-3* (*chitinase*), and *PR-5* (*thaumatin-like protein*), the phenylpropanoid pathway gene *PAL* (*phenylalanine ammonia-lyase*) and genes involved in plant stress responses and innate immunity, such as *LOX1* (*lipoxygenase*) and *NtPrxN1* (*peroxidase*), were strongly induced until 120 h after pGM spray application. Accumulation of superoxide radicals was also observed in a pGM dose-dependent manner.

## Introduction

To face pathogen attack, plants have developed important pathways of response against infections, which include mainly the immunity associated with the recognition of conserved pathogen or microbe molecular patterns (PAMPs/MAMPs), such as fungal chitin or bacterial flagellin by plant cell receptors (PRRs) (reviewed by e.g. [1]). This first line of response is called PAMP-triggered immunity (PTI). To overcome PTI, pathogens develop effector molecules that enable them to successfully infect a host even when PTI is active. Plants, however, can face this by a second line of defense, effector-triggered immunity (ETI) [2, 3]. PTI may result in a decrease in pathogen colonization, consequently blocking the disease development and conferring basal resistance. Second, ETI confers resistance later, often resulting in a hypersensitive response (HR). Immunity directly associated with pathogen effectors is triggered by host cell identification of the effector itself or by effector-induced responses. This recognition accelerates and amplifies PTI-mediated responses, often leading to an HR at infection sites and to a systemic induction of resistance known as systemic acquired resistance (SAR). Once properly stimulated, SAR provides long-term defense against a broad spectrum of pathogens [4–8]. Therefore, a localized microbial infection in a single or in some leaves can immunize the rest of the plant against a subsequent infection. This phenomenon makes the plant temporally resistant to new infection events even if the subsequent infection occurs at a site far from the initial primary site.

*Cladosporium herbarum* is a fungus that is not pathogenic to humans, but the easy dispersion of its spores in air makes it an important airway allergen, leading to the development of respiratory diseases such as rhinitis, conjunctivitis and asthma [9]. In addition, it has also been described as a phytopathogen in some agricultural crops, such as passion fruit and corn. *C. herbarum* infects passion fruit, causing cladosporiosis, which reduces fruit production and quality [10]. In corn, it causes *Cladosporium* ear rot [11]. Cladosporiosis can occur by contaminated seedlings or wind-dispersed conidia [12], potentially affecting any aerial parts of the plant but mainly growing tissues such as leaves, branches, flower buds and fruits, negatively impacting plant development and production [13].

The cell walls of members of the *Cladosporium* genus have a complex composition consisting mainly of polysaccharides (80-90%) and, to a lesser extent, proteins, glycoproteins and lipids [14]. Chitin and β-glucans are the main polysaccharides and are located in the inner layer of the cell wall, whereas glycoproteins are anchored in the outermost layer [15–17]. Fungal cell wall composition may vary according to the morphological type, growth stage and fungal species and is different from the plant cell wall composition, which mainly includes cellulose [18]. Among the fungal cell wall glycoproteins, peptidogalactomannan (pGM) can be found in various fungi, such as *Cladosporium werneckii, C. resinae, Aspergillus fumigatus, Aspergillus wentii, Malassezia* species and *Chaetosartorya chrysella* [19–22].

In a previous study, the *Cladosporium herbarum* pGM structure was elucidated and found to include carbohydrates (76%) and mannose, galactose and glucose as its main monosaccharides (52:36:12 molar ratio). The presence of a main chain containing (1→6)-linked α-D-Man*p* residues was observed by methylation and ^13^C-nuclear magnetic resonance (^13^C-NMR) spectroscopy [23]. β-D-Galactofuranosyl residues were present as (1→5)-interlinked side chains of *C. herbarum* pGM. The role of pGM cell wall glycoprotein in plant-fungus interactions was first studied by Mattos *et al*. [23], who showed the induction of the expression of defense genes in tobacco BY2 cells and a hypersensitive response (HR) after treatment of tobacco leaves with *C. herbarum* pGM. This recognition and defense response activation would possibly contribute to the plant response to the pathogen attack, protecting itself against new infections and indicating that pGM is probably able to induce SAR.

Here, using two distinct *Nicotiana tabacum* cultivars, Xanthi and SR1, we evaluated the ability of *C. herbarum* pGM to induce antiviral activity against *Tobacco mosaic virus* (TMV), PR genes and oxygen-reactive species (ROS) accumulation. Our results showed that pGM treatment induced a SAR-like response in both tobacco cultivars with strong activation of defense responses by the treated plants, enabling them to impair the viral infection.

## Material and Methods

### Strain and culture conditions

*Cladosporium herbarum*, CBS 121621, was provided by Dr J. Guarro, Advanced Studies Institut, Réus, Spain and was maintained in potato dextrose broth (PDB/Acumedia) in Erlenmeyer flasks at room temperature for 7 days with shaking. Mycelium was obtained via filtration, washed with distilled water and stored at −20°C.

### Extraction of *C. herbarum* glycoprotein

Crude glycoprotein extraction was performed according to [22]. Briefly, *C. herbarum* mycelium was extracted with 0.05 M phosphate buffer, pH 7.2, at 100°C for 2 h. The mixture was filtered, and the filtrate was evaporated into a small volume and precipitated with three volumes of ethanol overnight at 4°C. The precipitate was dialyzed and freeze-dried to obtain the crude glycoprotein (pGM). pGM purity was checked by HPTLC and GC-MS as described by Mattos *et al*., [23] and no contaminants were present.

### Plant growth and pGM treatment

*N. tabacum* cv. Xanthi and cv. SR1 seeds were germinated and grown in substrate in a greenhouse at 25 ± 2°C with a natural photoperiod. Young adult tobacco plants with 4-6 true leaves were sprayed with 600 μg.ml^−1^ of water diluted pGM with a high-pressure apparatus (W550, Wagner) according to [24]. MilliQ water was sprayed as control. Four pGM assay experiments were performed with Xanthi cv. with 16 plants sprayed with pGM and mechanically infected with TMV 24 h later and 16 plants sprayed with water and mechanically infected with TMV 24 h later in each experiment. Plants without treatment were used as healthy controls, plants only mechanically inoculated with TMV were used as TMV inoculation controls, plants treated with pGM were used as pGM control plants, and plants just sprayed with water were used as water treated (H_2_O) controls (mock). pGM assays with SR1 cv. were carried out 3 times with 10 plants used for each control (healthy plants, pGM alone, H_2_O-mock and TMV alone) and 15 plants for pGM + TMV and 15 for H_2_O + TMV in each experiment. Immediately before spraying, pGM was resuspended in MilliQ water, and the suspension was sterilized by filtration in a Millipore 0.20 μm filter.

### *Tobacco mosaic virus* (TMV) infection and evaluation of disease severity

For TMV mechanical infection, 1 g of TMV-infected frozen leaves (−80° C) was ground with a mortar and pestle in 19 ml of 0.01 M potassium phosphate buffer pH 7.0 to obtain the viral suspension [25]. Twenty-four hours after water or pGM treatment, the first 3 true leaves of each plant were mechanically inoculated with TMV without the use of abrasive. Experiments were carried out 4x with *N. tabacum* Xanthi and 3x with *N. tabacum* SR1 plants.

For cv. Xanthi, 48 hours post infection (hpi) with TMV, the disease incidence was measured by direct counting of necrotic lesions in each infected leaf, as described by [26], where the number of necrotic lesions was expressed as the infection percentage.

To assess TMV disease severity in pGM treated cv. SR1 plants, the plants were evaluated by visual observation 25 days post TMV infection (dpi) using a 0-5 disease scale. In this scale, the value 0 corresponds to the absence of symptoms; 1 to one or two leaves showing light mottling; 2 to more than two leaves showing light mottling with few thin yellow veins; 3 to mottling and vein clearing unevenly distributed on the leaf; 4 to mottling, leaf distortion, and stunting; and 5 to severe mottling, leaf curling, and stunting. The severity of the disease was quantified using the disease index (DI%) proposed by [27], applying the following formula: *DI* = ∑(*DS* × *P*)/(*TNP* × *HGS*) × 100, where DS = degree of the scale determined for each plant; P = number of plants showing each degree of infection (score); TNP = total number of plants evaluated; and HGS = highest grade of the scale (maximum infection score).

### ELISA assays

Enzyme-Linked Immunosorbent Assays (ELISA) analyses were carried out to measure the amount of the virus in leaf samples using PathoScreen^®^ (Agdia) ELISA kit for specific detection of tobamovirus family including TMV following Agdia protocol. Leaf samples of water- and pGM-treated SR1 plants inoculated with TMV were collected 72 h after virus infection. Each sample was ground in a 1:10 (leaf:extraction buffer (Adgia Co.) dilution and then loaded onto ELISA 96-well plates. Tests were performed in technical triplicates and the plates were read using a microplate spectrophotometer from BioRad Co.. Virus quantification was obtained using the standard curve of a 1.0 x 10^7^ TMV particles/g of leaf sample serial dilution.

### Measurement of superoxide radical accumulation in leaves treated with pGM

Evaluation of superoxide radical accumulation was performed using nitroblue tetrazolium (NBT) staining according to [28]. Leaves sprayed with 100, 200, 400 and 600 μg.ml^−1^ of pGM were collected after 24 and 72 h and 8 and 10 days after spray application and incubated with NBT at 0.5 mg.ml^−1^ for 1 h in vacuum. Subsequently, leaves were immersed in 95% boiling ethanol until for total removal of chlorophyll.

### Analysis of defense gene expression by qRT-PCR

Total RNA was extracted from young leaves between 24 and 120 h after treatment with water or 600 μg.ml^−1^ pGM using TRIzol^®^ Reagent (AmbionRNA by Life Technologies™) according to the manufacturer’s instructions. RNA concentration and purity were determined using a NanoDrop2000 Spectrophotometer (Thermo Scientific Co.). RNA integrity was assessed using 1% agarose gel electrophoresis and ethidium bromide staining. One microgram of total RNA from each sample was treated with RQ1 RNase-Free DNase (Promega Co.) according to the manufacturer’s instructions. Complementary DNA (cDNA) synthesis was performed with a Revert Aid First Strand cDNA Synthesis Kit (Fermentas Co.) and 100 μM of OligodT primer using 1 μg of total RNA as a template, according to the manufacturer’s instructions. Following cDNA synthesis, the samples were diluted 25-fold in sterile water. Primers for qPCR, described by [23], were designed for seven defense-related genes (*PR-1a*, *PR-2*, *PR-3*, *PR-5*, *NtPrxN1*, *LOX1* and *NtPAL*) and two constitutively expressed genes (*PP2A* and *Nt-ACT9*) Amplification reactions were performed on an Applied Biosystems^®^ 7500 Fast Real-Time PCR apparatus using a 96-well plate. All reactions were performed using two independent biological pools composed of leaves of 5 independent plants each. Three technical triplicates were analyzed for each biological replication. Three negative controls without cDNA were included on the plate for each primer. A mix was performed according to the manufacturer’s instructions containing the specific primer pairs for each gene at 10 μM and SYBR Green/ROX qPCR Master Mix (Thermo Scientific). cDNA amplification reactions were performed in a final volume of 25 μl, according to the manufacturer’s guidelines. qPCR cycles were 10 minutes at 95°C for initial denaturation, followed by 40 cycles of denaturation at 95°C for 15 sec and annealing/extension at 60°C for 1 minute, except for the PAL and PR5 genes, for which the annealing temperature was adjusted to 62°C [23]. The results were analyzed by the 2^−ΔΔ^ CT method according to [29].

### Statistical analysis

Statistical analyses were performed using GraphPad Prism software version 5.00 for Windows using “ONE-WAY ANOVA” Bonferroni test to evaluate if total number of TMV-induced necrotic lesions differ significantly between pGM treated and water-treated plants and compare ELISA results between treatments. Disease severity index between treatments were compared using One Way ANOVA Kruskal-Wallis and Dunn’s Multiple Comparison Test and “TWO-WAY ANOVA” Bonferroni test was used to evaluate qRT-PCR results.

## Results

### pGM induces tolerance against TMV infection in tobacco plants

To evaluate the potential of *C. herbarum* pGM in inducing virus defense in *N. tabacum*, tobacco plants from two distinct cultivars were sprayed with 600 μg.ml-1 pGM and mechanically inoculated with TMV 24 hours later. Forty-eight hours after TMV infection, plants from the tobacco cultivar Xanthi sprayed with water showed TMV-induced typical necrotic lesions (Fig 1A middle panel). Plants sprayed with pGM, however, showed a reduction in the number of necrotic lesions after TMV infection. A reduction of 42% in the number of necrotic lesions was observed when comparing water- and pGM-treated plants after TMV challenge and of 51% comparing pGM-treated and untreated TMV-infected plants (TMV infection control) (Fig 1B). Statistical analyses, however, showed that the number of necrotic lesions of water-treated and untreated plants were similar. Seven days after TMV infection, leaves at position 3 of plants previously sprayed with water showed 2.6x more necrotic lesions than leaves at the same position of pGM-sprayed plants (Fig 1C and 1D).

**Figure 1:**
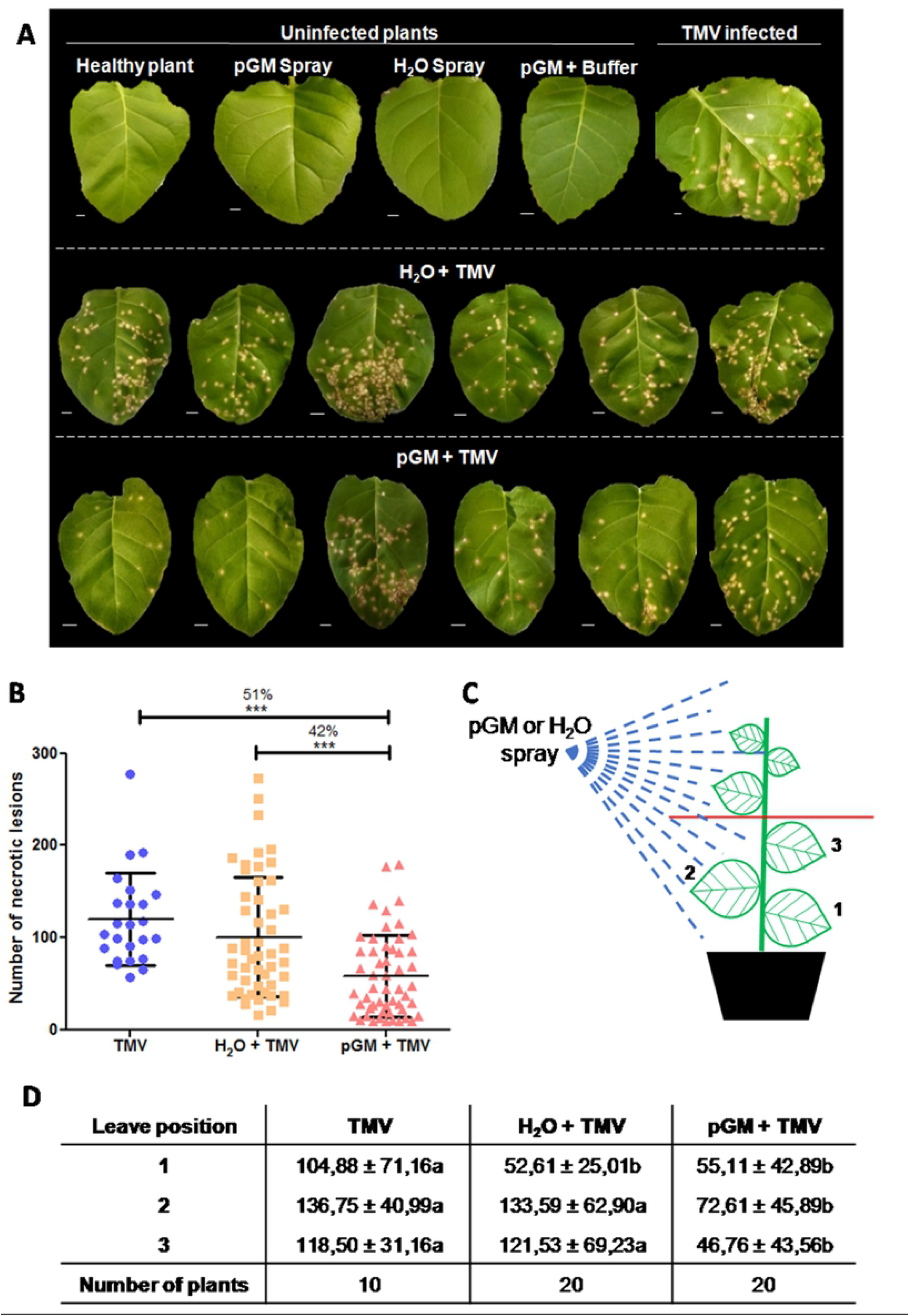
pGM treatment induced a reduction in the number of TMV induced necrotic lesions in *Nicotiana tabacum* cv. Xanthi. *N. tabacum* cv. Xanthi were treated with water or pGM and mechanically inoculated with TMV 24 hours later. (A) Details of representative leaves 7 dpi with TMV. Upper panel shows TMV infected untreated plants. Middle and bottom panels show leaves from water and pGM-treated plants, respectively. (B) Number of necrotic lesions in pGM treated plants. Dot plots represent the sum of the necrotic lesions observed in leaves 1-3 of each plant after TMV infection in untreated (n=10), H_2_O (n=20) and pGM (n=20) treated plants. Horizontal bars represent average values and vertical bars SE. The percentage of necrotic lesions reduction is shown over the dots and *** indicates significant differences with *p* < 0.001. (C) pGM treatment experimental scheme. After pulverization, leaves 1-3 were infected with TMV. (D) Total number of TMV-induced necrotic lesions at leaves 1-3 after each treatment from a representative experiment. Different letters represent statistical differences between the treatments for each leaf position with a *p* value < 0.05. Experiments were repeated two times. pGM - peptidogalactomannan. TMV - *Tobacco mosaic virus*. Bar: 1 cm

Tobacco plants from the TMV-susceptible cv. SR1 were similarly assayed. Symptoms of TMV infection were evaluated 3 weeks after infection. Typical symptoms of tobacco mosaic disease were visible since 18 dpi on untreated control as well as on water-treated plants. A strong reduction in disease symptoms was observed in pGM-treated plants, where only mild symptoms were observed. Fig 2A shows representative leaves of pGM treated infected plants. A disease severity index was used to compare TMV disease severity of individual plants after each treatment. As shown in Table 1, the pGM treatment induced a decrease between 76-80% in the disease severity compared to treatment with water, suggesting that pre-treatment with pGM could confer protection to the plants against TMV infection. Elisa assays performed in the infected plants showed that viral accumulation decreased 10 times on pGM-treated TMV infected SR1 plants compared to water-treated TMV infected plants (Table 1 and Fig 2B).

**Figure 2:**
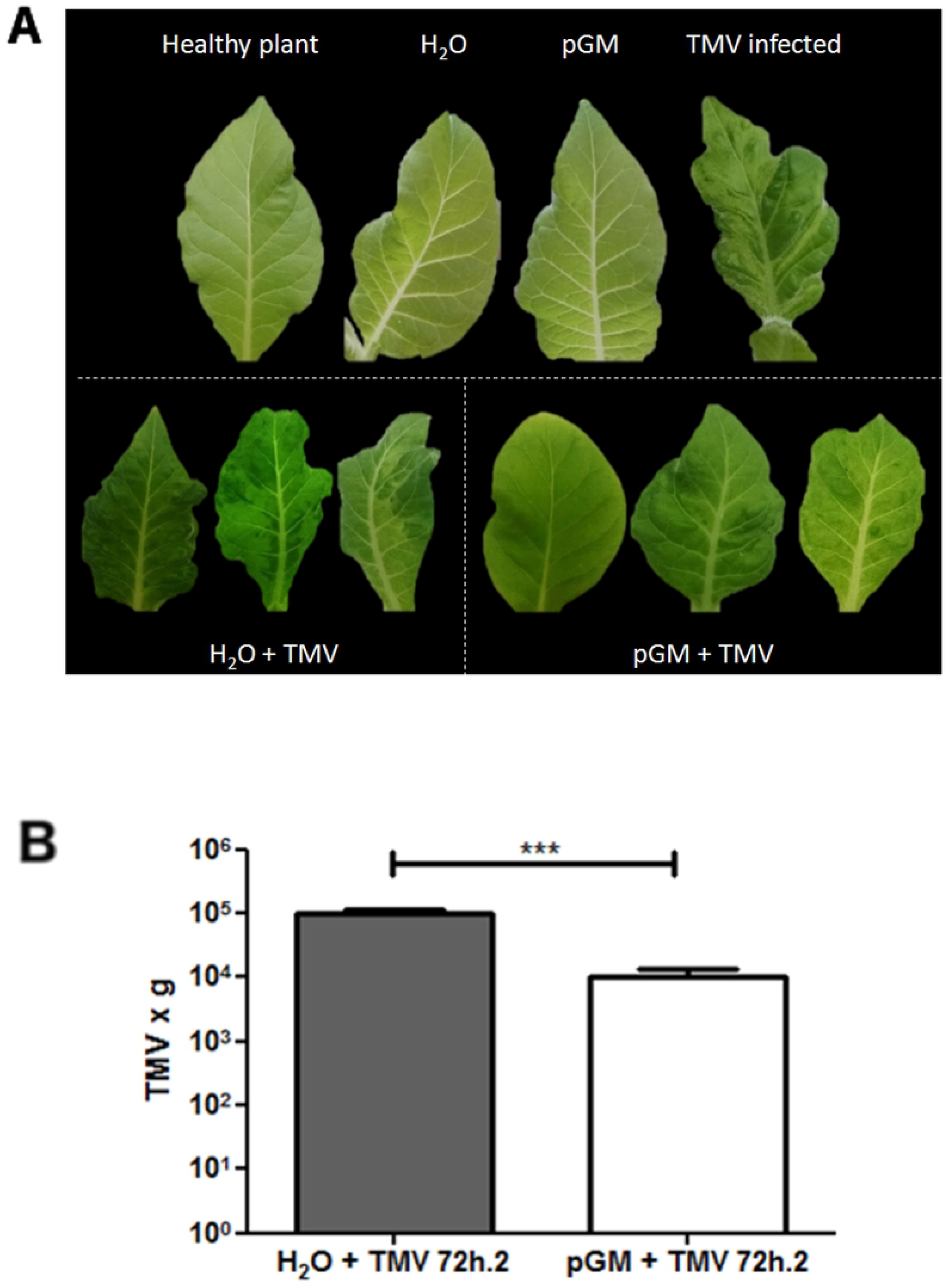
pGM treatment induced TMV tolerance in *N. tabacum* cv. SR1. *N. tabacum* cv. SR1 were treated with water or pGM and mechanically inoculated with TMV after 24 h. (A) Leaves of controls (upper panel) and water and pGM-treated TMV infected plants 22 dpi. (B) TMV accumulation seventy two hours post TMV inoculation. Levels of TMV particles were assayed in water and pGM treated plants by ELISA assay. ODs observed in water and pGM treated plants were plotted in a TMV quantification standard curve. Boxes are showing the average number of TMV particles per g of leaf. Vertical bars are showing SE and the horizontal bar shows ANOVA Bonferroni test comparing viral accumulation on water and pGM treatment with *p* < 0.01.

**Table 1:** Response of tobacco SR1 pGM treated plants to tobacco mosaic virus (TMV).

| Treatments | Number of symptomatic plants*<br>Experiment 1 | Disease severity index (%)**<br>Experiment 1 | Number of symptomatic plants*<br>Experiment 2 | Disease severity index (%)**<br>Experiment 2 | TMV detection by ELISA (415 nm)<sup>§</sup> | TMV quantification |
| --- | --- | --- | --- | --- | --- | --- |
| untreated | 10/10 a | 100.0 a | 10/10 a | 100.0 a | $1.70 \pm 0.14$ a | $1.0 \times 10^5$ a |
| H<sub>2</sub>O | 20/20 a | 100.0 a | 20/20 a | 100.0 a | $1.29 \pm 0.03$ a | $2.0 \times 10^5$ a |
| pGM | 12/20 b | 20.0 b | 14/20 b | 24.0 b | $0.35 \pm 0.17$ b | $1.0 \times 10^4$ b |

Number of TMV symptomatic per infected plants (\*) and a disease severity index (\*\*) shown by *N. tabacum* cv. SR1 25 dpi with TMV for the distinct treatments from two independent experiments.
§ - ODs average and SE obtained by TMV ELISA detection at 415 nm.
Column 7 shows the average number of TMV particles per g of infected leaf in each treatment.
Different letters correspond to significant differences with $p < 0.001$ . Statistical analysis was performed using One way ANOVA Bonferroni test (columns 2-4 and 6-7) and Kruskal-Wallis and Dunn's Multiple Comparison Test (columns 3 and 5).

So, pGM treatment was able to induce virus tolerance in both tobacco cvs., with a reduction in the number of necrotic lesions in Xanthi and an important decrease of mosaic disease in SR1.

### ROS accumulation in pGM-treated tobacco plants

It is proposed that ROS can act as defense compounds against viruses [30, 31]. H_2_O_2_ can act as a systemic antiviral signaling molecule during TMV infection; however, the role of ROS in plant–virus interactions is not completely understood [32]. The accumulation of ROS is also considered a biochemical marker of SAR induction [31].

To check if pGM treatment induces ROS accumulation in the tobacco treated plants, we examined the presence of ROS in the pGM-sprayed plants. Leaves from plants sprayed with different pGM concentrations (100, 200, 400 and 600 μg.ml^−1^) were analyzed over time, and the presence of superoxide radicals was assayed using nitroblue tetrazolium (NBT) reduction and histological staining. Figure 3 illustrates the accumulation of superoxide radicals after treatment with different pGM concentrations. We observed a dose-dependent effect of pGM spray on superoxide radical accumulation. Superoxide accumulation was observed from 24 h to 10 days after pGM spray for all pGM concentrations analyzed. After 8 days, however, a decay in its accumulation was observed. Curiously, 400 μg.ml^−1^ pGM showed a stronger induction of superoxide radical accumulation than 600 μg.ml^−1^, showing that the capacity of ROS induction of pGM may be saturated at higher concentrations. Histological assays to detect the presence of hydrogen peroxide in these plants using 3,3-diaminobenzidine (DAB) were also performed; however, DAB deposition was not observed (data not shown).

**Figure 3:**
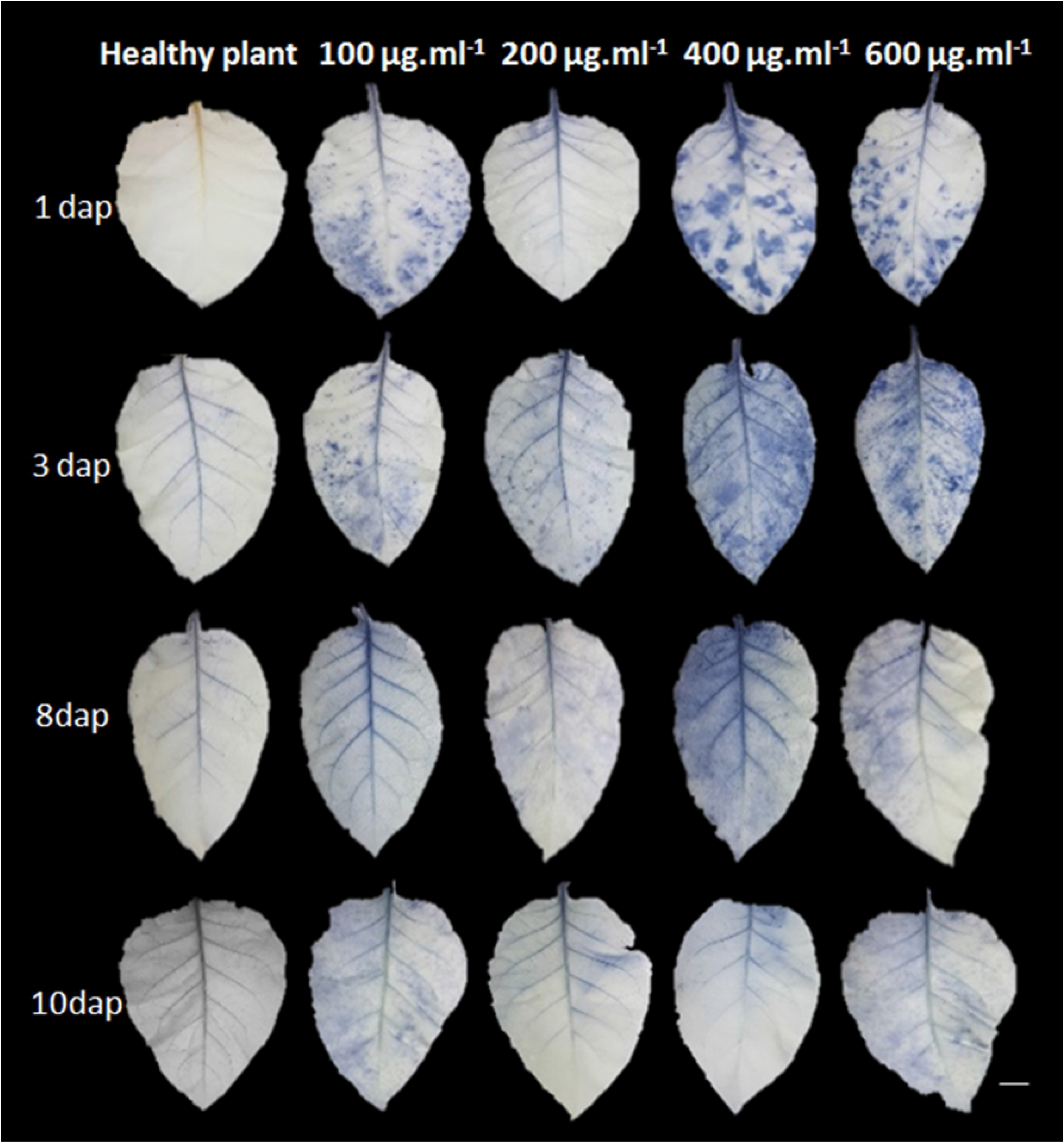
pGM is inducing superoxide radical accumulation in pGM-treated Xanthi plants. The accumulation of superoxide radicals after spray of pGM at different concentrations (100, 200, 400 and 600 μg.ml^−1^) was evaluated along 10 days using nitroblue tetrazolium (NBT). Representative leaves of each treatment are shown. dap - days after pGM pulverization. Bar: 1cm.

### pGM treatment induces defense gene expression in tobacco plants

To analyze whether pGM spray induces defense-related gene expression in treated plants, the transcript levels of the *pathogen-related 1-3* and *5* (*PR1α, PR2, PR3* and *PR5*), as well as *peroxidase* N1 (*NtPrxN1*), *phenylalanine amnonia-lyase* (*PAL*) and *lypoxigenase* 1 (*LOX1*) genes, were evaluated over time in *N. tabacum plants* from cvs. Xanthi and SR1 after spray application of 600 μg.ml^−1^ pGM.

pGM induced the expression of all PR genes analyzed 24 h after treatment in both tobacco cvs (Fig 4). A very strong induction of the PR1-α expression, a classical SAR marker, was observed in Xanthi. Twenty-four hours after pGM spray, *PR1*-α expression was more than 2000-fold compared to that in control plants sprayed with water. However, this gene expression decreased 24 h later and increased again 72 h after pGM treatment but at lower levels than during the first 24 h. Looking at the *PR1-α* expression in cv. SR1, we observed a more pronounced expression at 72 h, reaching approximately 300-fold that of the control. An up- and down-regulation cycle was observed in plants from this cv., where we observed an increase of more than 30x in *PR1-α* expression in the first 24 h, followed by a small decrease of 4x at 48 h and again a drastic increase at 72 h. After that, *PR1-α* expression levels decreased again at 96 h and returned to high levels at 120 h. In summary, *PR1-α* was strongly induced by pGM treatment in both tobacco cultivars, however, in Xanthi, the induction and its decay occurred earlier than in SR1. The presence of high levels of *PR1* transcripts in SR1 pGM-treated plants after 96 and 120 h after pGM spray may indicate that pGM treatment is in fact inducing systemic spreading signals over the sprayed leaves as the leaves samples of these two points developed after pGM treatment.

**Figure 4:**
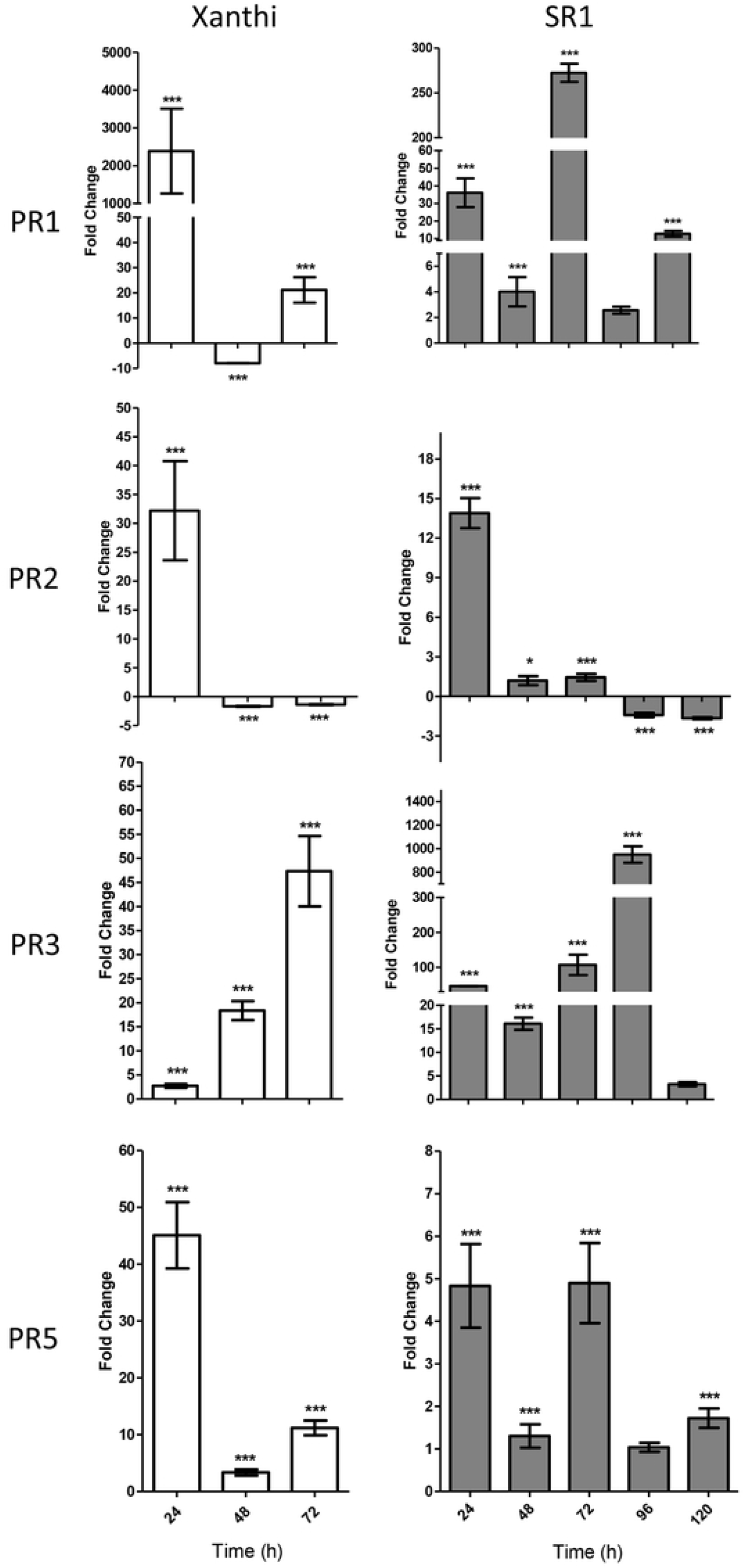
qRT-PCR analysis of some PR genes in water and pGM-treated plants between 24 and 120 h after treatment. Axis y is showing the fold change of *PR1-α, PR2, PR3* and *PR5* transcripts levels in *N. tabacum* cv. Xanthi and cv. SR1 after pGM treatment in comparison with water treatment calculated using 2^−ΔΔCt^ method described by [29]. *PP2A* and *NtACT-9* were used as reference genes. The standard deviations are indicated by error bars, and significant differences between fold changes with *p* < 0.05 and *p* < 0.001 are indicated by * and ***, respectively. *PR1-α*: Unknown function, possible antifungal function; *PR2*: *β- (1,3) endonuclease*; *PR3*: *chitinase*; *PR5*: *thaumatin-like protein*. *PP2A* and *NtACT-9* were used as reference genes.

Xanthi and SR1 showed 30- and 14-fold increases in *PR-2* (*β-1,3 glucanase*) gene expression after 24 h, respectively (Fig 4). After this time, however, *PR-2* mRNA levels were little reduced in water- and pGM-treated Xanthi plants and little induced in SR1, showing an early induction only of this specific gene. *PR-3*, which encodes the *chitinase* gene, showed a very low induction (3x) in Xanthi in the first 24 h but an increase of 50x in SR1 plants at the same time. With increasing time, an elevation in its expression was observed, reaching more than 45-fold that of the control at 72 h in Xanthi. SR1 *PR-3* induction mediated by pGM was stronger and peaked at 96 h after pGM treatment (more than 900-fold change expression) and decreased with time. Expression of *PR-5*, which encodes a *thaumatin-like protein* gene, was 45- and 5-times higher than that of the control after 24 h in Xanthi and SR1 plants, respectively, decreasing with time in Xanthi.

*NtPrxN1* (*peroxidase*) transcripts were highly induced in Xanthi as well in SR1, reaching values more than 200x at 24 and 72 h and 110x at 72 h in Xanthi and SR1, respectively (Fig 5). The high expression of the *peroxidase* gene may indicate that abnormal levels of ROS accumulated after pGM treatment in both cultivars. Leaves that developed after pGM spray, however, did not show *NtPrxN1* mRNAs induction (Fig 5, 96 and 120 h).

**Figure 5:**
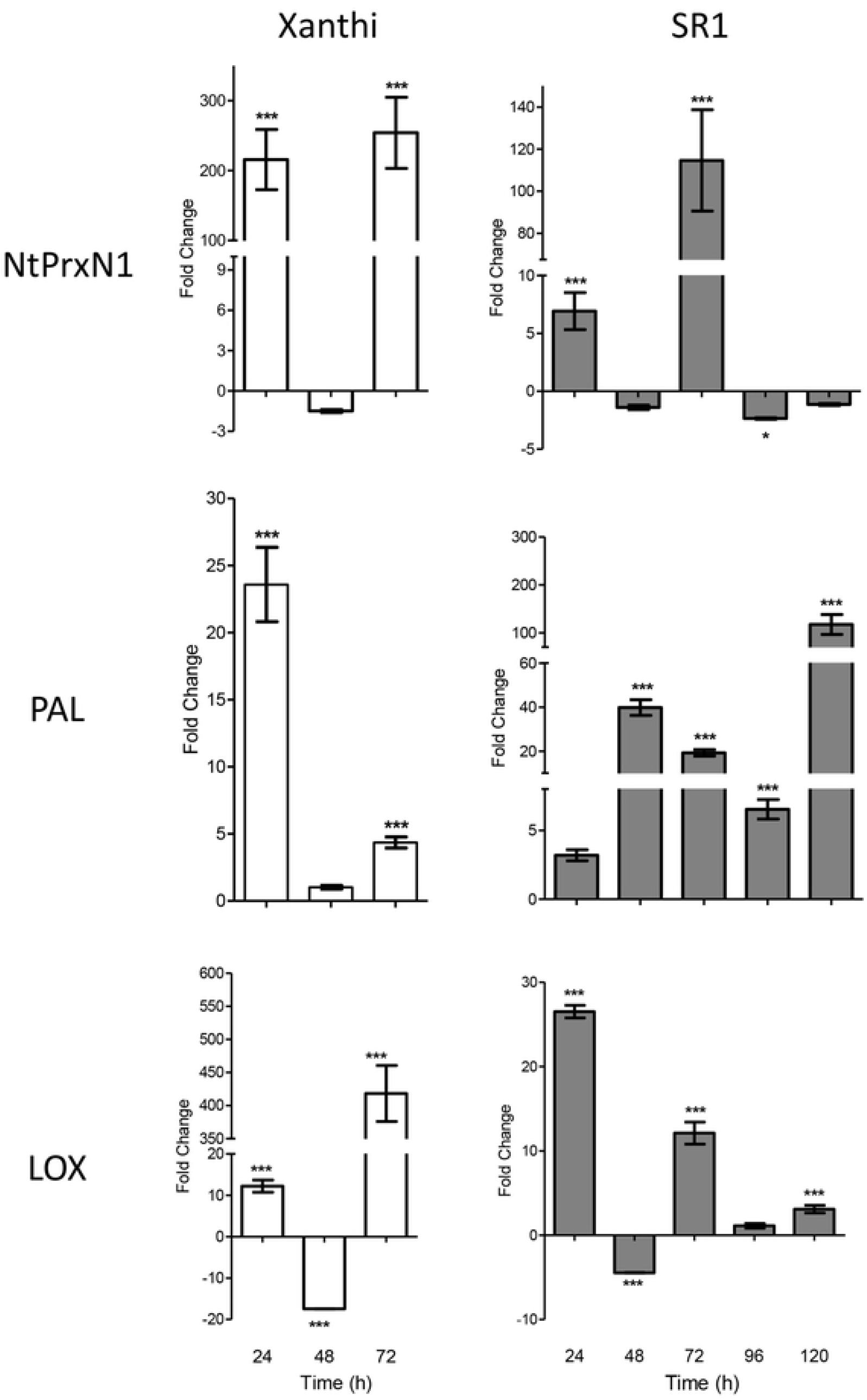
qRT-PCR analysis of defense related genes in water and pGM-treated plants between 24 and 120 hours after treatment. Axis y is showing average fold changes between transcripts levels of *NtPrxN1* (*peroxidase*), *PAL* (*phenylalanine amnonia-lyase*) and *LOX* (*lipoxygenase*) of *N. tabacum* cv. Xanthi and cv. SR1 after pGM treatment in comparison with water treatment calculated using 2^−ΔΔCt^ method described by [29]. *PP2A* and *NtACT-9* were used as reference genes. The standard deviations are indicated by error bars, and significant differences between fold changes with *p* < 0.05 and *p* < 0.001 are indicated by * and ***, respectively.

The *PAL* (*phenylalanine ammonia-lyase*) gene was also induced in both tobacco cultivars; however, the time course of *PAL* expression was different between them. In Xanthi, expression was induced earlier, reaching its highest level at 24 h after pGM spray application (24x). After this time point, the expression decreased to levels similar to those of the control. In SR1, however, the expression was only 5x more than that of the control at 24 h, reaching a maximum at 120 h, where it was expressed 130-fold compared with the control. *LOX* (*lipoxygenase*) gene, also associated with oxidative stress, was strongly induced (425x) in Xanthi plants at 72 h after pGM treatment. In SR1 plants, a comparatively slighter expression (27x) of this gene was observed, peaking at 24 h after pGM treatment.

Our RT-qPCR assays showed that almost all of the *PR*s and defense-related genes analyzed were strongly induced by pGM treatment. Our results indicated that pGM is responsible for the activation of a local systemic defense, observed in samples collected between 24-72 h. Results obtained from SR1 leaves that do not receive pGM directly (collected at 96 and 120 h), suggest that pGM may be an SAR inducer.

## Discussion

The present work aims to unravel the role of the *C. herbarum* cell wall peptidogalactomannan (pGM) as a plant defense elicitor. Treatment of young *Nicotiana tabacum* cv. Xanthi plants with pGM before TMV infection caused a reduction of 51 and 42% in the number of TMV-induced necrotic lesions when compared with untreated plants and water-sprayed plants, respectively. In *N. tabacum* cv. SR1 plants, decrease of 76-80% in disease severity was detected after treatment with *C. herbarum* pGM, showing that pGM treatment is able to promote protection against the viral pathogen in both tobacco cvs. In agreement, ELISA assays confirmed a reduction in the viral accumulation after treatment. In addition, we also observed a dose-dependent accumulation of superoxide radicals in pGM-treated plants. Superoxide radical accumulation seemed to be especially high during the first 24 h after pGM treatment, decreasing after 10 days. It was interesting to observe that even doses lower than 600 μg.ml^−1^, used in this work, and 400 μg.ml^−1^, previously used with BY2 tobacco cells [23], were able to induce the accumulation of superoxide radicals, indicating that oxidative stress mediated by ROS is induced in leaves after pGM treatment. A significant strong increase in expression of *NtPR1-α* transcript levels was also observed after treatment. The *PR1-α* gene is an SAR marker associated with salicylic acid (SA) and SAR signaling. Our data showed that in Xanthi as well as in SR1 tobacco plants, *PR1-α* transcript levels were highly induced, suggesting local defense and possible SAR were induced by pGM. Importantly, pGM treatment did not induce any damage to the fitness of the plants. In contrast, treated plants even showed slight growth enhancement compared to water- and untreated plants (data not shown).

Recently, our group showed that treatment of tobacco roots and BY2 cells with *C. herbarum* pGM induces oxidative stress and the expression of *PR1-α* and other defense-related genes [23]. In addition, an HR-like response was observed when pGM was infiltrated in tobacco leaves. To our knowledge, our results show for the first time that a peptidogalactomannan isolated from the *C. herbarum* cell wall is able to induce local and systemic defense responses in a whole plant. In the literature, other reports have shown that distinct glycoproteins are able to confer SAR protection to a plant. Baillieul *et al*. [33] and Cordelier *et al*. [34], using a 32 kDa glycoprotein named alpha-elicitin secreted by the oomycete *Phytophthora megasperma*, demonstrated the induction of the HR and the production of enzymes related to SAR after its administration in tobacco leaf mesophyll. A reduction in the size of TMV-induced necrotic lesions in leaves was observed after treatment with the glycoprotein. These studies, however, did not show a reduction in the number of necrotic leaves, as observed in our work. In addition to the reduction in the number of necrotic lesions observed on the HR TMV-resistant tobacco Xanthi, we also observed a reduction in protection mediated by pGM in TMV-susceptible SR1 cv., with a strong reduction in disease severity. Glucan 1,4-alpha-glucosidase (BcGs1), isolated and purified from *Botrytis cinerea* culture supernatant, was also able to induce SAR in tomato and tobacco plants [35]. Enzyme-treated plants showed the induction of necrotic lesions that mimicked a typical HR. H_2_O_2_ production was also increased in the treated tomato and tobacco plants that exhibited resistance to *B. cinerea*, *Pseudomonas syringae* pv. tomato DC3000 and *Tobacco mosaic virus* along with an increase in the transcript levels of the defense-related genes *PR1-α*, *TPK1b* and prosystemin and a reduction of approximately 40% in TMV-induced necrotic lesions.

Several authors also reported the use of bacterial, fungal and oomycete culture filtrates and/or secreted molecules to induce plant defense responses. The protein PemG1, an elicitor molecule isolated from *Magnaporthe grisea* culture medium and expressed in an *E. coli* heterologous system, induced resistance to bacterial pathogens in rice and *Arabidopsis* [36]. Treatment with PemG1 did not inhibit bacterial growth but increased plant resistance, indicating that PemG1 is an SAR elicitor. LI *et al*. [37] described a novel HR-inducing protein elicitor, called PeFOC1, isolated from the culture filtrate of *Fusarium oxysporum* f. sp. *cubense*. This protein induced ROS and an HR in tobacco cells and induced the expression of *PR* genes (with upregulation of *PAL, EDS1*, *LOX* and *PDF*). The SA and JA/ET signaling pathways were activated with the consequent induction of callose and phenolic compound deposition, causing an immune response and SAR in tobacco. A study by KWAK and collaborators [38] showed that an uncharacterized fraction obtained from aqueous extraction of the fungus *Hericium erinaceus* induces defense genes and promotes plant growth and death of the bacteria that cause disease in tomatoes.

In our previous work, we observed that the expression of defense-related genes such as *PrxN1*, *PAL*, *LOX* and the pathogen-related genes *PR1-α*, *PR-2* and *PR-3* were strongly induced after the treatment of tobacco BY2 cells with pGM [23]. In the present work, we observed that treatment of both tobacco Xanthi and SR1 with pGM also induced the expression of *PR-1α, PR-2, PR-3, PR-5, LOX1, PAL* and *NtPrxN1*. The strong induction of *PR1-α* and *PR-2* transcripts after pGM treatment suggests a possible SAR activation by pGM. Despite small differences in the expression levels in Xanthi and SR1, we observed that defense response induction mediated by pGM persisted for at least 3 and 5 days based on *PR1-α* and *PR-2* qRT-PCR results and the NBT assay.

Overexpression of *PR1-α* as well as other defense-related genes seems to directly contribute to elicitor-mediated pathogen resistance, as already demonstrated by [39]. The ability of PR genes to enhance resistance against both biotic and abiotic stresses is well documented. A strong induction of more than 2000x and 35x of the *PR1-α* gene during the first 24 hours after pGM treatment was detected in both Xanthi and SR1 tobacco plants, respectively, showing that the expression of the *PR1-α* SAR marker is induced in both tobacco plant cultivars. It is also interesting to highlight the increased expression of transcripts from the hydrolytic □-1,3-endoglucanases (*PR-2*), commonly activated in a fungal infection and another classical SAR marker, by treatment with pGM. Activation of this gene is directly related to an antimicrobial effect due to hydrolysis of □-1-3 glucans of the fungal wall, altering its integrity. Like □-1-3 endoglucanase, mRNAs of PR-3 or chitinase, another well-known antifungal protein, were strongly upregulated after 72 h of pGM treatment in both Xanthi and SR1 cv. SINDELAROVA & SINDECOR [40] reported that both *PR-2a* and *PR-3* from *Nicotiana tabacum* showed strong antiviral activity against TMV. In addition, peroxidase and PAL transcripts, which were also strongly induced in our treated plants, have also been shown to present antiviral activity [32, 41]. PAL may be induced by ROS accumulation and is also considered an SAR component and a key enzyme in defense, leading to the synthesis of antimicrobial molecules, including phytoalexins and pathogen-related proteins and to the strengthening of the physical barrier against pathogen colonization by deposition of structural polymers, such as lignin and callose, at the infection site [42, 43]. The upregulation of *PR1-α, PR-2, PR-3, peroxidase* and *PAL* genes observed after pGM treatment may explain the antiviral effects observed in our TMV infection assays for both Xanthi and SR1 tobacco plants. Interestingly, Xanthi and SR1 plants, however, showed differences in the time when these genes reached their peak of induction. Apparently, the NN Xanthi cv. recognizes the fungal elicitor faster than plants from SR1 cv.. Besides that, in Xanthi, the level of expression of these genes were also higher than in SR1, with exception to *PR-3* and *PAL*. In combination, these data suggest that Xanthi may be more prepared to respond against the pGM PAMP/MAMP than SR1. However further experiments would be necessary to test this hypothesis. As well, genome-wide transcriptome sequencing of pGM treated and untreated samples would provide a better insight into the genes underlying pGM-mediated defense response.

## Conclusions

The results obtained in this work allow us to suggest that *C. herbarum* pGM is a fungus PAMP able to induce the expression of *PRs* and *PAL*, *LOX* and *NtPrxN1* and to activate other classical SAR markers, such as ROS induction and protection against infection. Due to the importance of PR proteins in biotic and even abiotic stress tolerance, several researchers are trying to obtain multi tolerant transgenic plants by individual or dual PR overexpression (reviewed by [44]). Our data show that treatment with pGM may induce at least four *PR* genes simultaneously that help plants resist pathogen attack.

## Author contributions

Conceptualization: CBM, MFSV and EB-B. Formal analysis: CBM, BBM, MFSV and EB-B. Funding acquisition: MFSV and EB-B. Investigation: CBM. Methodology: CBM, TFS. Project administration: MFSV and EB-B. Resources: MFSV and EB-B. Supervision: MFSV and EB-B. Writing – original draft: CBM, MFSV and EB-B. Writing – review & editing: CBM, TFS, EB-B and MFSV.

## Conflict of interest

The authors declare that they have no conflict of interest.

